# From 3D Time-of-Flight Angiography to Accelerated 4D Arterial Spin Labeling Angiography: A Fast Few-Shot Transfer Learning Approach

**DOI:** 10.64898/2026.05.18.725892

**Authors:** Hao Li, Iulius Dragonu, Peter Jezzard, Thomas W. Okell, Mark Chiew

## Abstract

**Purpose:** To develop a data-efficient deep learning framework for rapid reconstruction of highly accelerated 4D arterial spin labeling (ASL) magnetic resonance angiography (MRA) with robust generalization using extremely limited acquired data, addressing the challenges of prolonged acquisition and reconstruction time.

**Methods:** A simulation-driven, few-shot transfer learning approach was adopted by leveraging publicly available 3D time-of-flight (TOF)-MRA data to generate realistic multi-coil complex-valued pseudo-ASL k-space datasets for large-scale pre-training. A 3D unrolled reconstruction network was trained on this simulated data using a histogram-weighted loss and subsequently extended to 4D using lightweight temporal fusion modules. Fine-tuning was performed using only two experimentally acquired 4D ASL-MRA datasets. The method was evaluated on retrospectively and prospectively undersampled Cartesian 4D ASL-MRA data acquired at 3T and compared with compressed sensing (CS) and locally low-rank (LLR) reconstructions.

**Results:** The proposed method achieved superior reconstruction quality compared with CS and LLR, with improved vessel depiction, particularly in distal branches, and enhanced temporal fidelity. Quantitative evaluation demonstrated higher vessel-masked peak signal-to-noise ratio and structural similarity index measure, along with increased error entropy, indicating reduced noise and structured artifacts. The initial pre-trained model already outperformed conventional methods, while additional 4D fine-tuning further improved performance. Robust reconstruction was demonstrated in prospectively undersampled data and multi-slab acquisitions, enabling large-coverage, time-resolved angiography within clinically feasible scan times (4-6 min).

**Conclusions:** Simulation-driven pre-training combined with few-shot fine-tuning enables accurate and rapid reconstruction of highly accelerated 4D ASL-MRA in data-limited settings. The proposed framework provides a practical pathway toward clinically feasible, non-contrast dynamic cerebrovascular imaging.

## 1 Introduction

Imaging of blood flow through the arterial vasculature is crucial for the diagnosis, treatment planning, and follow-up of a wide range of cerebrovascular diseases, including stenoses, occlusions, and vascular malformations. Magnetic resonance angiography (MRA) has become a key modality for this purpose, enabling non-invasive visualization of the intracranial arterial system without ionizing radiation or exogenous contrast agents.

Among available techniques, three-dimensional time-of-flight (3D TOF) MRA is widely used in clinical practice for high-resolution depiction of arterial anatomy.^1^ However, it provides only static information and does not capture the temporal evolution of blood flow. Time-resolved MRA techniques have therefore been developed, among which arterial spin labeling (ASL)^2–5^ enables non-contrast dynamic angiography (4D ASL-MRA) using magnetically labeled blood as an endogenous tracer. Through control-label subtraction, 4D ASL-MRA provides zero-background angiograms that enhance visualization of distal vessels and capture temporal flow dynamics, enabling assessment of flow alterations in cerebrovascular diseases such as Moyamoya disease.^6^ In addition, multiple studies have demonstrated the utility of 4D ASL-MRA in the evaluation of arteriovenous malformations and dural arteriovenous fistulas, where separation of arterial inflow and venous outflow is important.^7–12^

Despite these advantages, 4D ASL-MRA remains limited by long acquisition times, particularly when high spatial and temporal resolution are required. To accelerate 4D MRA, advanced reconstruction methods have been developed, often in combination with non-Cartesian acquisition trajectories such as stack-of-stars and 3D golden-angle radial sampling. Reconstruction techniques such as k-space-weighted image contrast (KWIC)^13,14^ exploit temporal redundancy by sharing k-space data across frames, but may introduce temporal blurring, particularly in regions of rapid flow. Compressed sensing (CS)–based approaches^15– 19^ enable acceleration by enforcing sparsity in appropriate transform domains, but they require long reconstruction times and careful hyper-parameter tuning and often exhibit degraded image quality at high acceleration factors. Low-rank and subspace methods^20–22^ further enhance acceleration by exploiting temporal correlations, representing the dynamic signal using a limited set of temporal basis functions. However, these approaches typically require the temporal dynamics to be represented within a predefined low-dimensional subspace, which may not fully capture complex flow patterns, particularly in smaller vessels. In addition, faster reconstruction remains desirable for routine clinical use.

Deep learning (DL)–based reconstruction has emerged as a promising alternative, offering rapid inference and improved reconstruction fidelity.^23^ However, its application to 4D ASL-MRA is highly constrained by the scarcity of large, fully sampled training datasets. Yousefi et al.^24^ have applied DL to jointly accelerate ASL angiography and perfusion imaging using a 3D convolutional neural network (CNN). To address data limitations, they augmented a small in-house ASL dataset through permutation and registration with anatomical images from the BrainWeb^25^ dataset. While this approach demonstrated promising results at relatively low spatial resolution and modest acceleration (R=2), its performance on high-resolution *in vivo* data remains untested. A recent study has employed CNNs for accelerated 4D ASL-MRA reconstruction^26^, demonstrating a substantial increase in reconstruction speed. However, due to the limited availability of fully sampled data, the method was trained using reference reconstructions derived from CS methods rather than true ground truth data, and relied on 2D networks that do not fully exploit through-slice information.

Fundamentally, the data scarcity poses a key barrier to the development of robust DL-based reconstruction methods for 4D ASL-MRA. Limited datasets prevent model generalization across populations, anatomical variability, acquisition protocols, and scanner platforms, while increasing susceptibility to overfitting and reducing reproducibility across sites.^23^ While self-supervised learning partially alleviates the need for fully sampled data by training directly on undersampled acquisitions, it still typically requires large volumes of k-space data to achieve stable performance.^27–29^ More recent zero-shot self-supervised methods^30^ further reduce reliance on training datasets, but depend on subject-specific optimization, leading to increased reconstruction times that hinder their clinical use.

To address these challenges, we propose a data-efficient deep learning framework for accelerated 4D ASL-MRA reconstruction. Building on our prior work in 3D TOF-MRA,^31,32^ which demonstrated that large-scale simulated k-space data can enable effective model pre-training, the present study extends this concept to dynamic ASL angiography. Specifically, widely available 3D TOF-MRA magnitude images are used to simulate diverse and realistic multi-coil 3D ASL k-space data for large-scale pre-training, followed by few-shot fine-tuning using only two acquired 4D ASL datasets. The pre-trained 3D reconstruction model is then efficiently extended to 4D through the incorporation of a temporal fusion module. The proposed method is quantitatively evaluated against established reconstruction approaches using retrospectively undersampled Cartesian 4D ASL-MRA *in vivo* datasets acquired from healthy subjects at 3T. In addition, prospectively undersampled acquisitions are used to assess robustness and temporal fidelity under real-world experimental conditions, and further validated through multi-slab acquisitions to demonstrate the potential for large-coverage dynamic angiography within clinically feasible scan and reconstruction times.

## 2 Methods

The proposed method consists of two main components: (i) simulation of realistic 3D multi-coil ASL k-space data from 3D TOF-MRA magnitude images for large-scale pre-training; and (ii) deep learning–based reconstruction using a physics-informed unrolled network extended from 3D to 4D with subsequent few-shot fine-tuning on limited *in vivo* 4D ASL-MRA datasets. An overview of the proposed simulation-driven, few-shot learning framework is shown in Figure 1. This builds upon our previous work presented in abstract form.^33^

**Figure 1.**
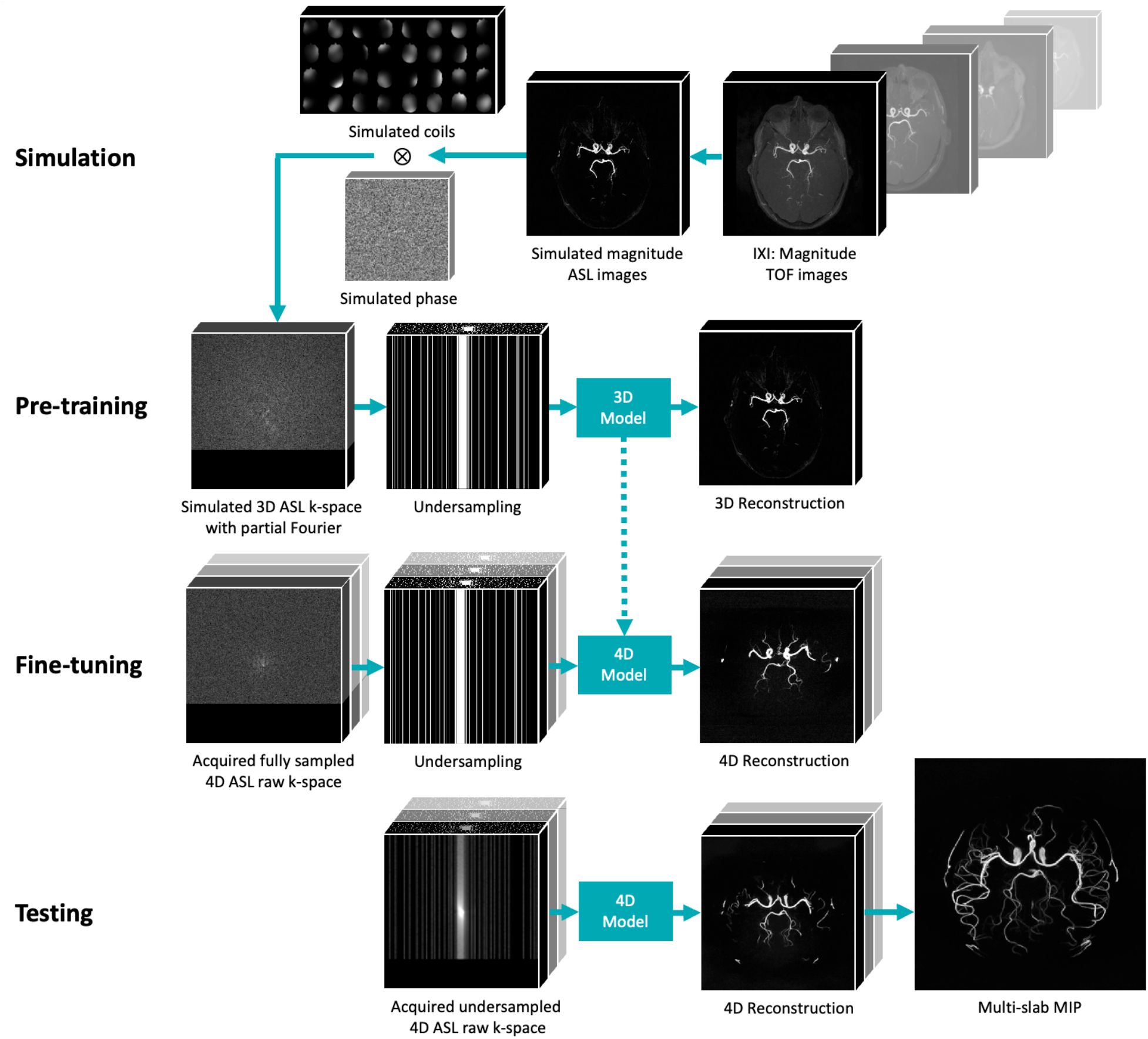
Proposed simulation-driven few-shot learning framework. Open-source 3D TOF-MRA magnitude images were used to simulate complex-valued, multi-coil 3D ASL-MRA k-space data after control–label subtraction. Poisson-disc undersampling was then applied retrospectively to pre-train a 3D DL reconstruction model. After pre-training, the model was extended from 3D to 4D and fine-tuned using only two experimentally acquired 4D ASL-MRA datasets on a 3T scanner. The final model was evaluated on additional 4D ASL-MRA datasets acquired independently. MIP: maximum intensity projection.

### 2.1 Data Simulation

To generate ASL-like training data from 3D TOF-MRA magnitude images from the large open-source multi-vendor multi-center IXI dataset (https://brain-development.org/ixi-dataset/), we developed a vessel-focused preprocessing pipeline designed to emphasize bright arterial signal while suppressing background structure, mimicking the removal of background tissue through the control-label subtraction process used in ASL. First, vessel signal was enhanced using separable 1D filters ([-1 0.5 1 0.5 -1]) ^34^ applied along each spatial axis, and the maximum value across directions was retained to emphasize vessel-like structures. The filtered images were then normalized to [0, 1] and subjected to gamma correction (power = 1.5) to increase vessel-to-background contrast. The background was further suppressed by clipping intensities below μ -1.5σ (μ and σ denote the image mean and standard deviation) followed by re-normalization. Finally, Gaussian noise with randomly sampled mean (0–0.1) and standard deviation (0.001–0.025) was added to introduce realistic variability across datasets. The small mean offset accounts for residual background signal arising from motion or system drift, as observed in experimental ASL data. As shown in Figure 2A, the simulated ASL magnitude images exhibit substantially reduced background signal and enhanced vascular structures compared to the original TOF images, closely resembling experimentally acquired ASL angiograms.

**Figure 2.**
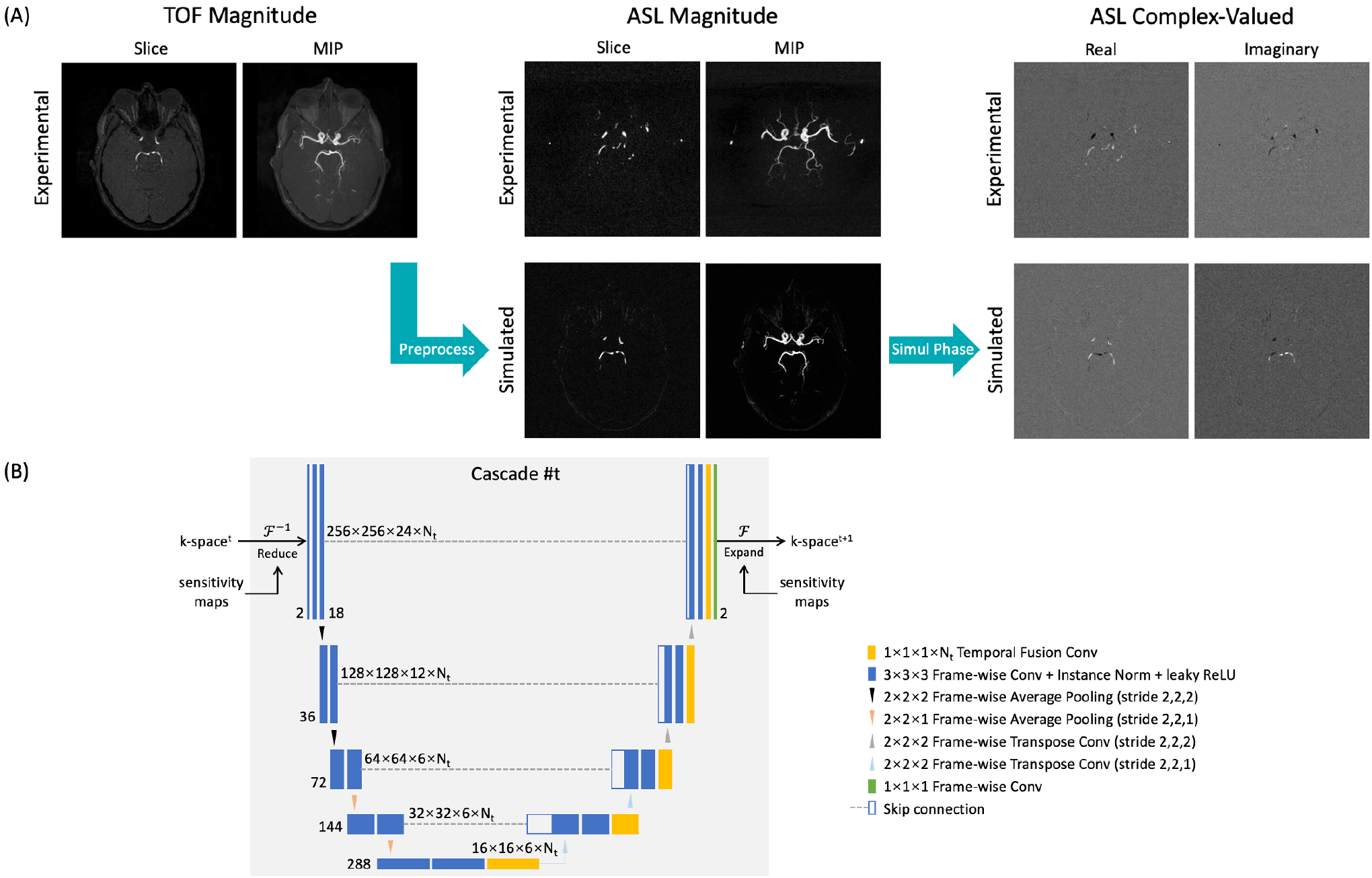
(A) Comparison of simulated and experimentally acquired ASL-MRA magnitude and complex-valued images. (B) Proposed 4D model architecture for each cascade in the unrolled reconstruction network. At the bottleneck and each decoder level, a 1×1×1×N_t_ convolution (where N_t_ is the number of temporal frames) was incorporated to fuse features across the time dimension.

Complex-valued ASL data were generated by simulating phase based on a structure-informed approach adapted from our previous work.^31^ Specifically, the conjugate-symmetric k-space of the magnitude images was perturbed with modulated noise components to introduce realistic phase variations. To further improve realism, we introduce region-specific perturbations to vessel and background regions, with parameter settings detailed in Table S1 (Supporting Information), yielding complex-valued images that closely match experimentally acquired ASL angiograms (Figure 2A). Multi-coil data were subsequently generated by applying simulated coil sensitivity maps using the same parameter settings as in the previous work.^31^

To further increase variability in the simulated data, spatially varying intensity bias fields were incorporated using the RandomBiasField transform from the TorchIO library (coefficients = 0.3, order = 3).^35^ Retrospective undersampling was then applied in the k_y_-k_z_ plane using 2D variable-density Poisson-disc masks with a fully sampled central calibration region (12 × 6). All simulation steps, except for undersampling, were performed once per dataset with randomly sampled parameters and then kept fixed throughout training to reduce computational and memory demands. For each training experiment, a fixed acceleration factor was used, while a new undersampling mask was randomly generated each time a sample was accessed, promoting robustness to variations in sampling patterns.

### 2.2 Deep Learning-Based Reconstruction

The pre-training stage employed a 3D unrolled neural network adapted from prior work^31^, consisting of cascaded data-consistency and regularization blocks. In each cascade, data consistency enforces agreement with the acquired k-space measurements using a forward model incorporating coil sensitivity maps, while the regularization is implemented using 3D U-Nets^36^ operating in the image domain.

In previous work, a L1+SSIM loss^37^ was used to balance pixel-wise fidelity and perceptual image quality. Here, to address the pronounced class imbalance between sparse high-intensity vascular structures and dominant background signal, we introduce a histogram-weighted L1+SSIM loss. Specifically, voxel-wise weights were assigned inversely proportional to the frequency of their intensity values, thereby emphasizing underrepresented vessel regions during optimization. This encourages improved reconstruction of fine vascular structures that are otherwise prone to attenuation or smoothing.

This 3D network was first pre-trained on large-scale simulated static ASL-MRA datasets generated from 3D TOF-MRA images, enabling the model to learn robust spatial representations of vascular anatomy and reconstruction priors without requiring extensive experimentally acquired ASL k-space data.

Following pre-training, the model was extended from 3D to 4D to incorporate temporal dynamics in ASL-MRA (Figure 2B). To preserve the spatial feature representations learned during pre-training, all convolutional kernels were retained and applied independently to each temporal frame. Temporal information was integrated through lightweight fusion modules implemented as 1×1×1×N_t_ convolutions at the network bottleneck and decoder levels, where N_t_ denotes the number of temporal frames. Notably, this design results in only a 13.8% increase in the number of trainable parameters, providing a practical and computationally efficient alternative to fully 4D spatiotemporal network architectures.

The pre-trained, extended 4D model was subsequently fine-tuned using only two fully sampled *in vivo* 4D ASL-MRA datasets. During fine-tuning, all network weights were updated, while the temporal fusion modules were randomly initialized. To prevent overfitting and improve generalizability, data augmentation was applied, including random flips along all spatial axes and random translations in the x-y plane (up to half the image size), with parameters resampled at each epoch. This few-shot transfer learning step allows the network to capture acquisition-specific characteristics and temporal signal evolution while retaining the generalizable spatial priors learned during pre-training. The overall framework thus enables data-efficient domain adaptation from 3D TOF-MRA in a setting with extremely limited 4D ASL datasets.

### 2.3 Training Data and *In Vivo* Acquisition

Simulated training data were generated from 3D TOF-MRA images of 341 subjects (153 males, 188 females; mean age 49 ± 17 years) from the publicly available IXI dataset (https://brain-development.org/ixi-dataset/). The dataset comprises multi-site acquisitions from Hammersmith Hospital, Guy’s Hospital, and the Institute of Psychiatry in London, United Kingdom, using 3T and 1.5T MRI systems from Philips Medical Systems (Best, The Netherlands) and GE HealthCare (Waukesha, WI, USA). Imaging parameters, obtained from the dataset documentation and NIfTI headers, included TR = 16.7 ms (3T) or 20 ms (1.5T), TE = 5.8 ms (3T) or 6.9 ms (1.5T), and flip angle = 16° (3T) or 25° (1.5T), with an acquisition matrix of 288 × 286, voxel size of 0.8 × 0.8 × 0.8 mm^3^, and 100 slices. To emulate multi-slab acquisition, each 3D magnitude volume was divided into five overlapping slabs along the slice direction, resulting in 24 slices per slab.

To evaluate the proposed DL-based reconstruction method on retrospectively undersampled data, a total of eight non-overlapping, fully sampled Cartesian 4D ASL-MRA datasets were acquired from five healthy volunteers (5 males; mean age 28 ± 2 years) on a 3T MAGNETOM Prisma scanner (Siemens Healthineers, Forchheim, Germany) with a 32-channel head coil, under a technical development protocol approved by local ethics and institutional review committees. A spoiled gradient-echo pseudo-continuous ASL (pCASL) sequence was employed, modified based on recent work^38^ to enable Cartesian acquisition. Cartesian sampling enables fully sampled ground-truth acquisitions for quantitative retrospective evaluation. Sequence parameters included TR/TE = 9.42/4.83 ms, quadratically varying flip angles^39^ = 4°–6°, acquisition matrix = 256 × 256 × 24, spatial resolution = 0.8 × 0.8 × 0.8 mm^3^, temporal resolution = 241 ms, number of frames = 4, and scan time =11 min 52 sec. Two datasets were used for fine-tuning (Cohort 1), and the remaining six datasets were reserved for testing (Cohort 2), with no subject overlap.

Prospectively undersampled datasets were then acquired using a modified version of the same sequence with predefined pseudo-random variable-density Poisson-disc undersampling masks, while keeping all other imaging parameters unchanged. Six non-overlapping datasets (Cohort 3) with 8× acceleration (1 min 29 sec each), along with six fully sampled reference datasets (11 min 52 sec each), were obtained from three healthy volunteers (2 males, 1 female; mean age 25 ± 2 years). In addition, two further undersampled datasets per subject were acquired to emulate non-overlapping multi-slab acquisitions for extended head coverage.

Finally, to demonstrate feasibility for large field-of-view dynamic angiography, a prospectively undersampled multi-slab 4D ASL-MRA acquisition was performed in one healthy volunteer. Imaging parameters included four slabs with 20% slab overlap, 20 slices per slab, and 20% slice oversampling. Accelerated datasets with factors of R=8 (6 min) and R=12 (4 min) were acquired. A 3D TOF-MRA dataset was also acquired with SENSE^40^ acceleration (R=2, offline reconstruction) as a reference using a vendor-provided sequence (TR = 21.0 ms, TE = 3.81 ms, flip angle = 18°) with matched matrix size, slabs and coverage and comparable scan time (5 min 4 s).

### 2.4 Experiments

Reconstruction performance was evaluated using both global and vessel-specific quantitative metrics. Peak signal-to-noise ratio (PSNR)^41^ and structural similarity index measure (SSIM)^42^ were computed to assess overall image fidelity. To specifically evaluate vascular structures, vessel-masked PSNR (VM-PSNR) and vessel-masked SSIM (VM-SSIM)^43^ were calculated using vessel masks derived from the fully sampled reference, following previous studies.^43,44^ These vessel-specific metrics emphasize angiographic regions, where background voxels otherwise dominate, and VM-SSIM has been shown to correlate strongly with radiologist assessments.^43,45^ All PSNR- and SSIM-based metrics were computed from axial maximum-intensity-projection (MIP) images. In addition, error entropy (EE)^46^ was computed from the reconstructed 4D volumes to quantify the randomness of residual errors, with higher values indicating more noise-like, unstructured artifacts. Together, these metrics capture both overall reconstruction fidelity and the preservation of vascular morphology, which is crucial for the clinical utility of 4D ASL-MRA.

Fully sampled reference images were reconstructed by combining multi-coil data using coil sensitivity maps estimated with ESPIRiT^47^, implemented via the BART toolbox.^48^ The proposed method was evaluated against these references and compared with spatiotemporally total variation– regularized CS,^49^ a locally low-rank (LLR)^50^ method recommended in previous work,^39^ and the pre-trained model without 4D fine-tuning (w/o 4D F-T). CS was implemented using the BART toolbox^48^ and LLR was implemented based on previously published code^51^. Hyper-parameters for CS and LLR were optimized on retrospectively undersampled datasets (Cohort 2) using vessel-focused metrics (VM-PSNR and VM-SSIM).

All models were pre-trained for 50 epochs using the Adam optimizer with a learning rate of 3 × 10^−4^, batch size of 3, and validation-based early stopping, minimizing the proposed histogram-weighted L1+SSIM loss. Pre-training of the 3D model required approximately 43 hours on an NVIDIA H100 80GB GPU. Fine-tuning was subsequently performed on two experimentally acquired 4D ASL-MRA datasets for up to 300 epochs with a batch size of 1, using early stopping based on two-fold cross-validation, requiring approximately 2.5 hours. Prior to reconstruction, 32-channel k-space data were compressed to 8 virtual coils using an SVD-based method^52^ to reduce computational cost. For prospectively undersampled datasets, reconstructed images were rigidly aligned to their corresponding fully sampled references using FLIRT^53,54^ from the FSL library (https://fsl.fmrib.ox.ac.uk/) to correct for inter-scan misalignment.

In addition, ablation studies were conducted on prospectively undersampled datasets (Cohort 3), comparing the proposed method with variants excluding the histogram-weighted L1+SSIM loss and excluding pre-training. To assess generalizability to altered temporal dynamics, the proposed method was also evaluated on time-reversed datasets. Furthermore, the impact of increasing the number of fine-tuning datasets was investigated by fine-tuning using both Cohort 1 and Cohort 2, with evaluation performed on Cohort 3.

## 3 Results

### 3.1 Retrospective Reconstructions

Representative reconstructions of retrospectively 8× accelerated single-slab 4D ASL-MRA data are shown in Figure 3 and Figure 4, with corresponding temporal MIP (tMIP) images. The reconstruction time of the proposed method was around 3.6 sec per dataset, compared with 158 sec for CS and 1,510 sec for LLR, corresponding to approximately 44× and 420× speedups, respectively.

**Figure 3.**
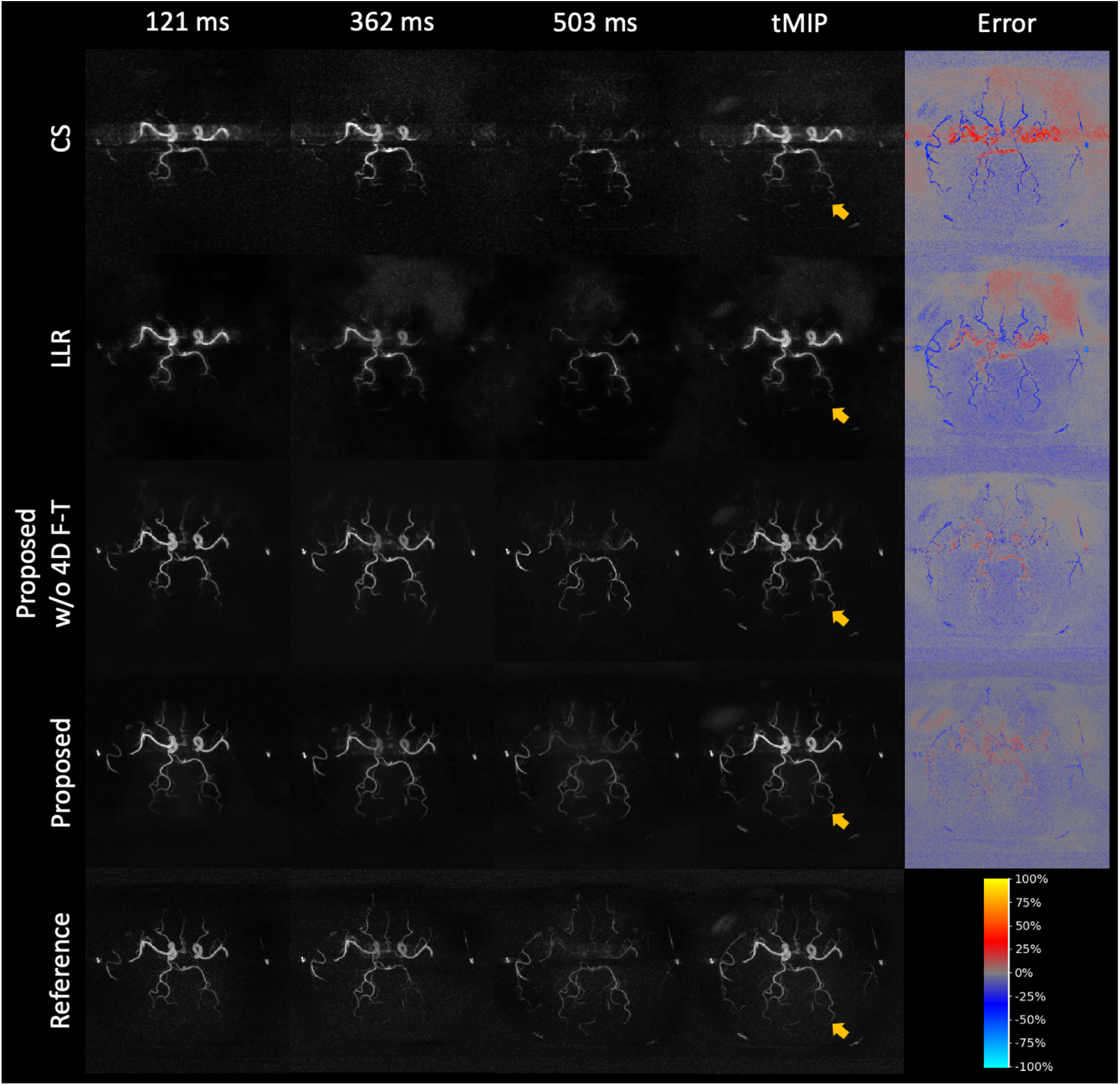
Axial MIP images of representative single-slab 8× retrospectively accelerated 4D ASL-MRA data reconstructed by CS, LLR, the proposed method without 4D fine-tuning (w/o 4D F-T), and the proposed method, alongside the fully sampled reference. The last column shows error maps (percentage difference from the reference normalized by the maximum intensity). Yellow arrows highlight representative differences between methods. tMIP: temporal MIP.

**Figure 4.**
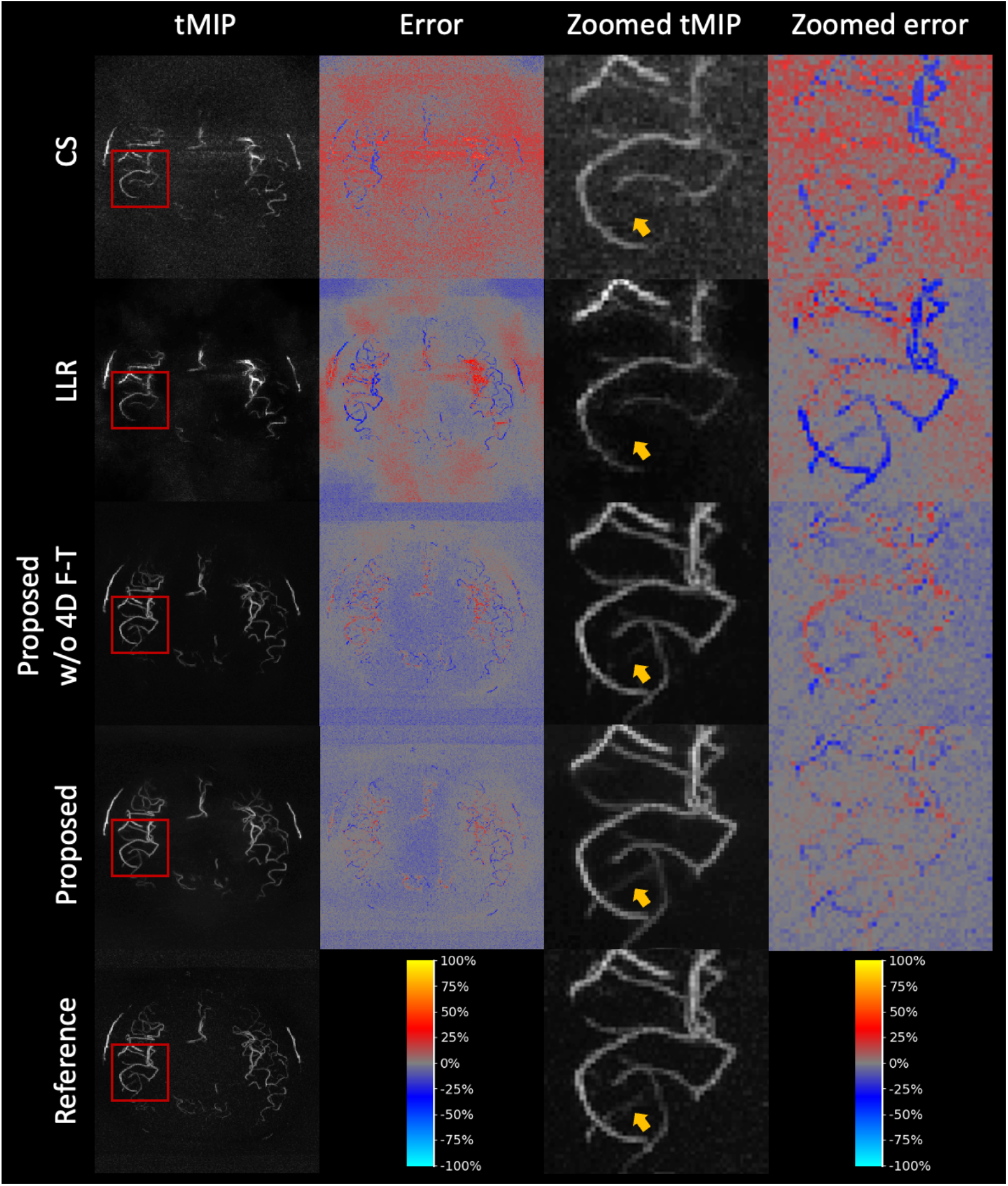
Temporal MIP (tMIP) images of another representative 8× retrospectively accelerated single-slab 4D ASL-MRA data acquired in a superior brain region reconstructed by methods as in Figure 3 with zoomed views.

Compared with CS and LLR, the proposed method consistently produced sharper vessel boundaries and improved continuity of distal branches. In Figure 3, CS-reconstructed images exhibit strong background noise and noticeable blurring around the middle cerebral artery (MCA), while LLR reduces noise but results in spatial smoothing and loss of fine vessels. The proposed method without 4D fine-tuning (w/o 4D F-T) partially recovers these structures but still shows attenuation of small vessels. In contrast, the proposed method reconstructs a more complete visualization of the vasculature with improved contrast and reduced artifacts. This is further reflected in the error maps, where structured residuals in vessel regions are substantially reduced.

Figure 4 highlights these differences in a superior brain region with more complex vascular branching. Zoomed views demonstrate that both CS and LLR fail to preserve small vessels, whereas the proposed method maintains their contrast and continuity. The improvement from 4D fine-tuning is also evident, where temporal information appears to stabilize reconstruction of weak vessel signals.

The proposed method also achieved the highest performance across all quantitative metrics (Table 1). Improvements were particularly evident in vessel-focused measures, with VM-PSNR increasing from 27.21 dB (CS) and 27.31 dB (LLR) to 33.38 dB, and VM-SSIM from 0.9384 (CS) and 0.9430 (LLR) to 0.9790, indicating improved reconstruction fidelity specifically within vascular regions. Similar trends were observed for global metrics, although their values are dominated by background signal. In addition, the proposed method achieved higher error entropy (3.387 vs 2.897 for CS and 2.920 for LLR), suggesting more noise-like and less structured residuals relative to the fully sampled reference.

**Table 1.** Quantitative comparison of reconstruction performance on 8× retrospectively accelerated 4D ASL-MRA data. Results presented as mean ± standard deviation across testing datasets. The top-performing method for each metric is highlighted in bold font. The proposed method showed significantly higher performance than all the comparison methods (p < 0.01).

| Metric | CS | LLR | Proposed w/o 4D F-T | Proposed |
| --- | --- | --- | --- | --- |
| PSNR ↑ | 21.06 ± 2.10 | 22.87 ± 0.44 | 24.60 ± 0.59 | **25.98 ± 0.68** |
| SSIM ↑ | 0.3172 ± 0.0734 | 0.3729 ± 0.0277 | 0.4664 ± 0.0318 | **0.5048 ± 0.0340** |
| VM-PSNR ↑ | 27.21 ± 1.43 | 27.31 ± 1.31 | 31.56 ± 1.24 | **33.38 ± 0.95** |
| VM-SSIM ↑ | 0.9384 ± 0.0065 | 0.9430 ± 0.0125 | 0.9643 ± 0.0167 | **0.9790 ± 0.0100** |
| EE ↑ | 2.897 ± 0.123 | 2.920 ± 0.134 | 3.280 ± 0.164 | **3.387 ± 0.168** |

Across temporal frames (Figure 5), the proposed method consistently outperformed other methods in both VM-PSNR and VM-SSIM, indicating improved preservation of both vascular details and temporal consistency.

**Figure 5.**
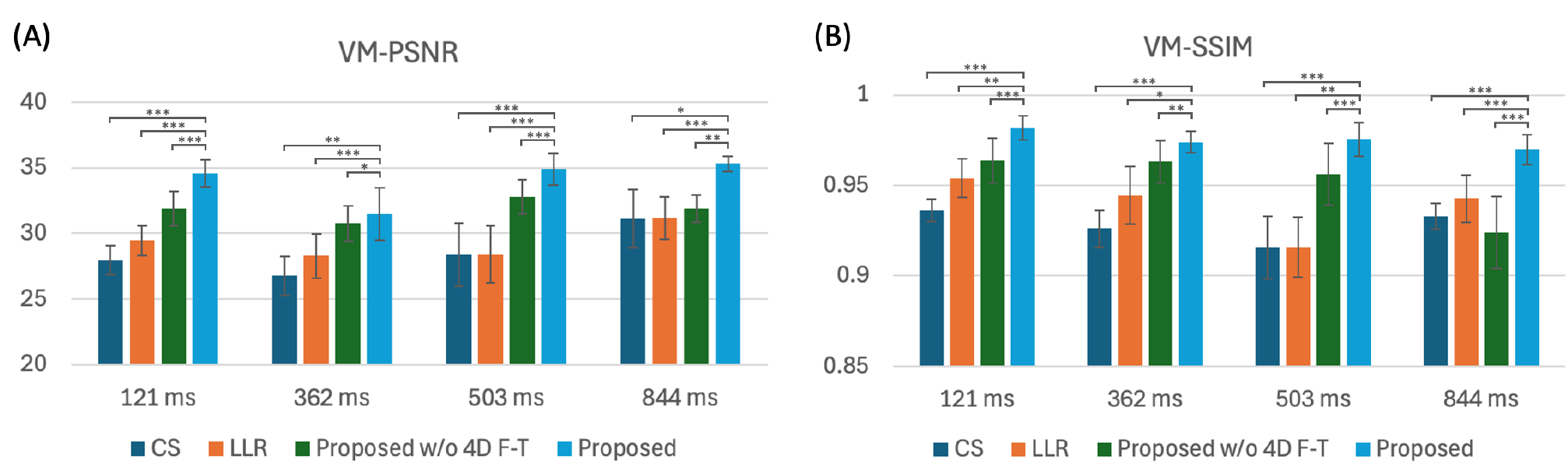
Quantitative comparison of reconstruction performance across temporal frames on retrospectively accelerated 4D ASL-MRA data. (A) VM-PSNR and (B) VM-SSIM evaluated at different temporal delays for CS, LLR, the proposed method without 4D fine-tuning (w/o 4D F-T), and the proposed method. Error bars indicate standard deviation across datasets. P values are indicated by stars: * P < 0.05, ** P < 0.01, *** P < 0.001.

### 3.2 Prospective Reconstructions

Results on prospectively accelerated datasets (Figure 6 and Figure 7) further demonstrate the superiority of the proposed method under realistic acquisition conditions. The proposed method consistently preserves vessel continuity while effectively suppressing noise, whereas CS and LLR exhibited increased blurring and residual artifacts. In Figure 7, zoomed tMIP images show that CS and LLR fail to recover fine vascular structures, particularly in regions affected by aliasing from bright MCA signals, whereas the proposed method preserves these structures with reduced error. Quantitative results in Table S2 (Supporting Information) further support these findings, with the proposed method achieving the highest performance across all evaluated metrics. Inter-scan misalignment, system drift, and physiological change between prospectively undersampled and fully sampled acquisitions likely contribute to overall higher errors compared with the retrospective study.

**Figure 6.**
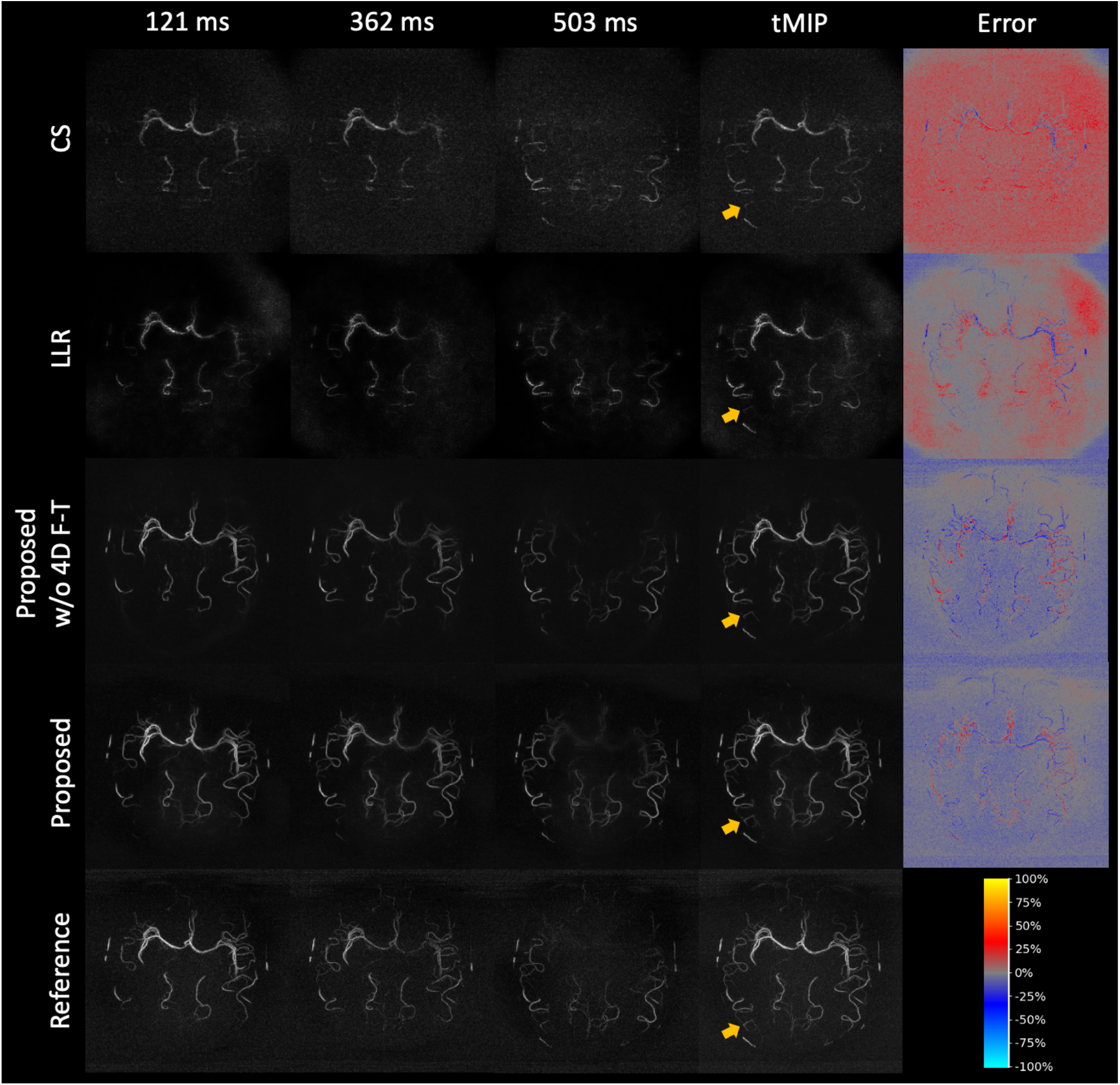
Axial MIP images of a representative single-slab 8× prospectively accelerated 4D-ASL-MRA dataset reconstructed by CS, LLR, the proposed method without 4D fine-tuning (w/o 4D F-T), and the proposed method, alongside the fully sampled reference. The figure layout follows Figure 3.

**Figure 7.**
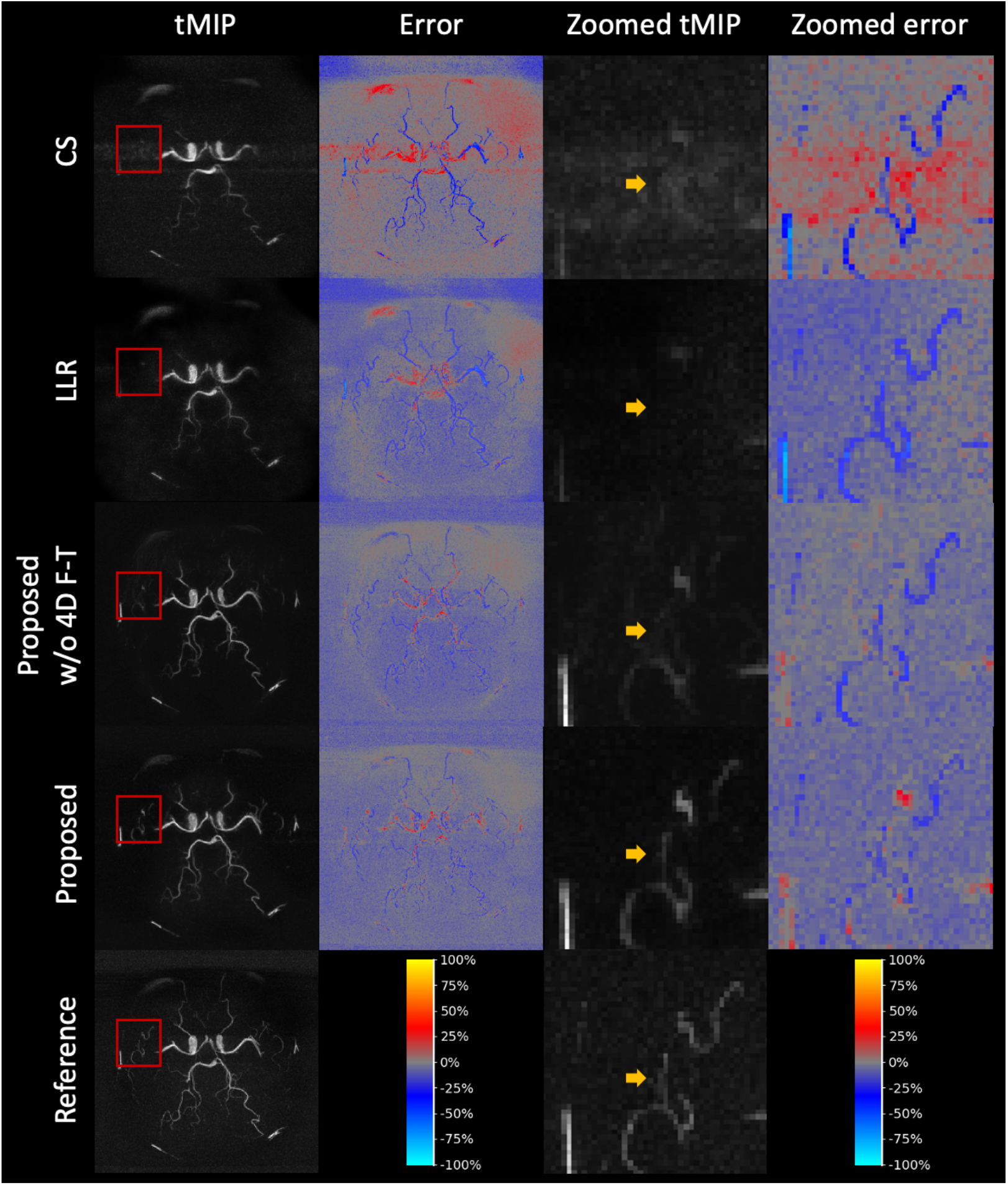
tMIP images of another representative 8× single-slab prospectively accelerated 4D-ASL-MRA data acquired in an inferior brain region reconstructed by methods as in Figure 6 with zoomed views.

Temporal signal fidelity is illustrated in Figure 8A. ASL signal curves extracted from selected regions of interest (ROIs; orange circles) show that the proposed method closely follows the fully sampled reference across all time points. In contrast, CS and LLR exhibit substantial deviations and inconsistent temporal trends. The model w/o 4D F-T shows larger deviations at peak inflow compared with the proposed method, confirming that 4D fine-tuning improves reconstruction of dynamic signal evolution.

**Figure 8.**
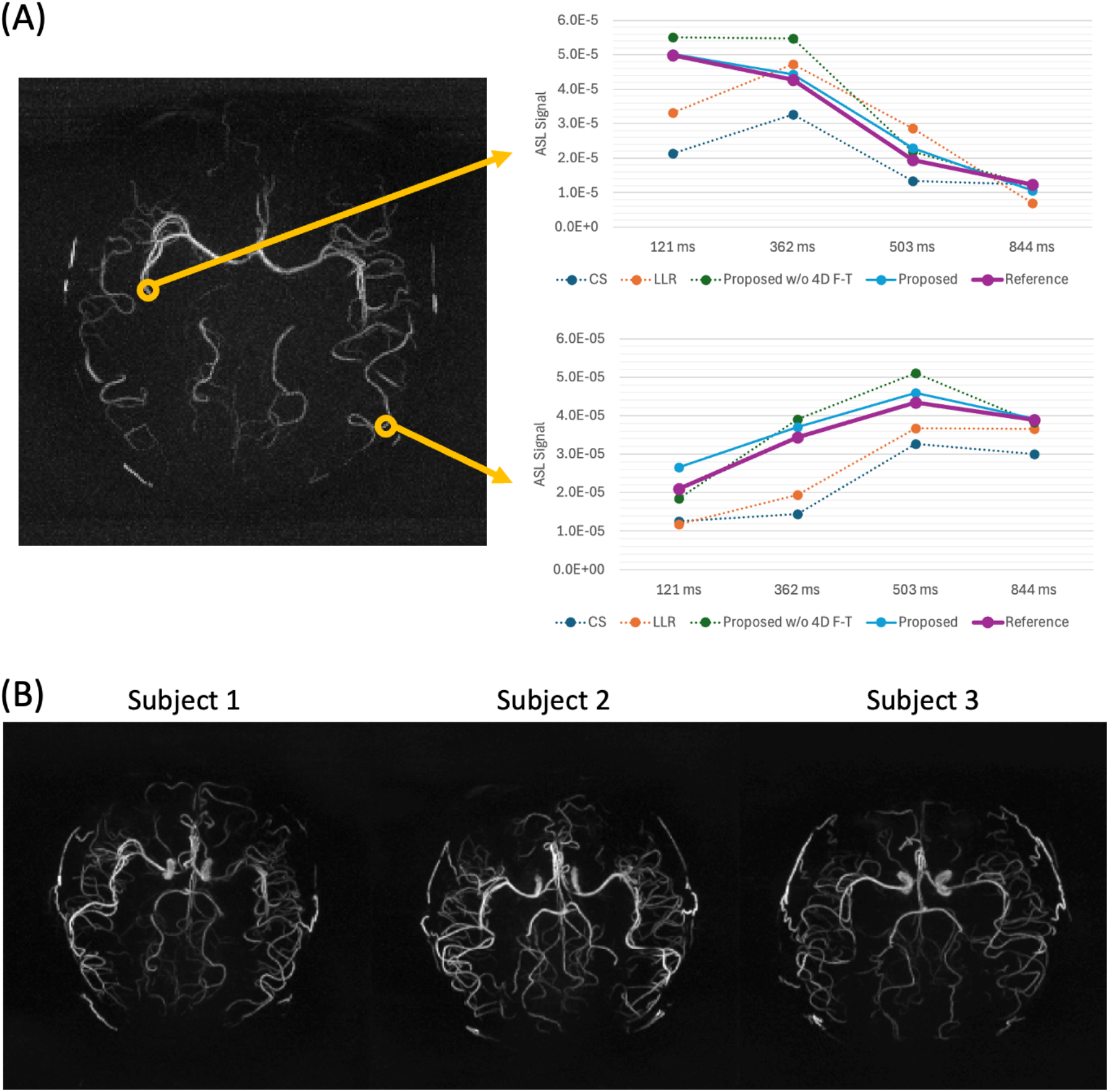
Temporal signal fidelity and multi-slab reconstruction for prospectively undersampled 4D ASL-MRA. (A) Representative MIP image with selected regions of interest (ROIs, orange markers) and corresponding ASL signal curves across temporal frames for methods shown in Figure 6. (B) Non-overlapping multi-slab axial MIP images reconstructed using the proposed method for three subjects.

Figure 8B demonstrates pseudo multi-slab reconstructions for three subjects using these non-overlapping prospectively undersampled datasets, showing consistent vascular coverage and continuity over extended fields of view. Due to the non-overlapping acquisition without oversampling, mild discontinuities and boundary artifacts are observed at slab interfaces (Figure S1, Supporting Information).

In addition, ablation study results (Table S3, Supporting Information) show that removing the histogram-weighted L1+SSIM loss or pre-training leads to consistently lower performance across all metrics, highlighting the importance of both components. Evaluation on time-reversed datasets resulted in a slight reduction in performance (Table S3), indicating that the proposed method remains reasonably robust to substantial changes in temporal signal evolution. The impact of increasing the number of fine-tuning datasets is shown in Figure S2 (Supporting Information), where performance improves substantially when increasing from zero to two datasets, with only marginal gains beyond this, suggesting diminishing returns with additional data.

To demonstrate the feasibility of rapid multi-slab 4D ASL-MRA acquisition for potential clinical use, Figure 9 compares multi-slab 2× accelerated 3D TOF-MRA (5 min 4 s) with prospectively undersampled 4D ASL-MRA at 8× (6 min) and 12× (4 min) acceleration. Despite the higher acceleration factors and additional dynamic information, ASL provides comparable anatomical information and even improved vessel visibility compared with TOF, particularly in distal arteries. At 12× acceleration, only minor degradation and slight noise amplification are observed, mainly affecting the weak vessel signals while preserving the main arterial tree.

**Figure 9.**
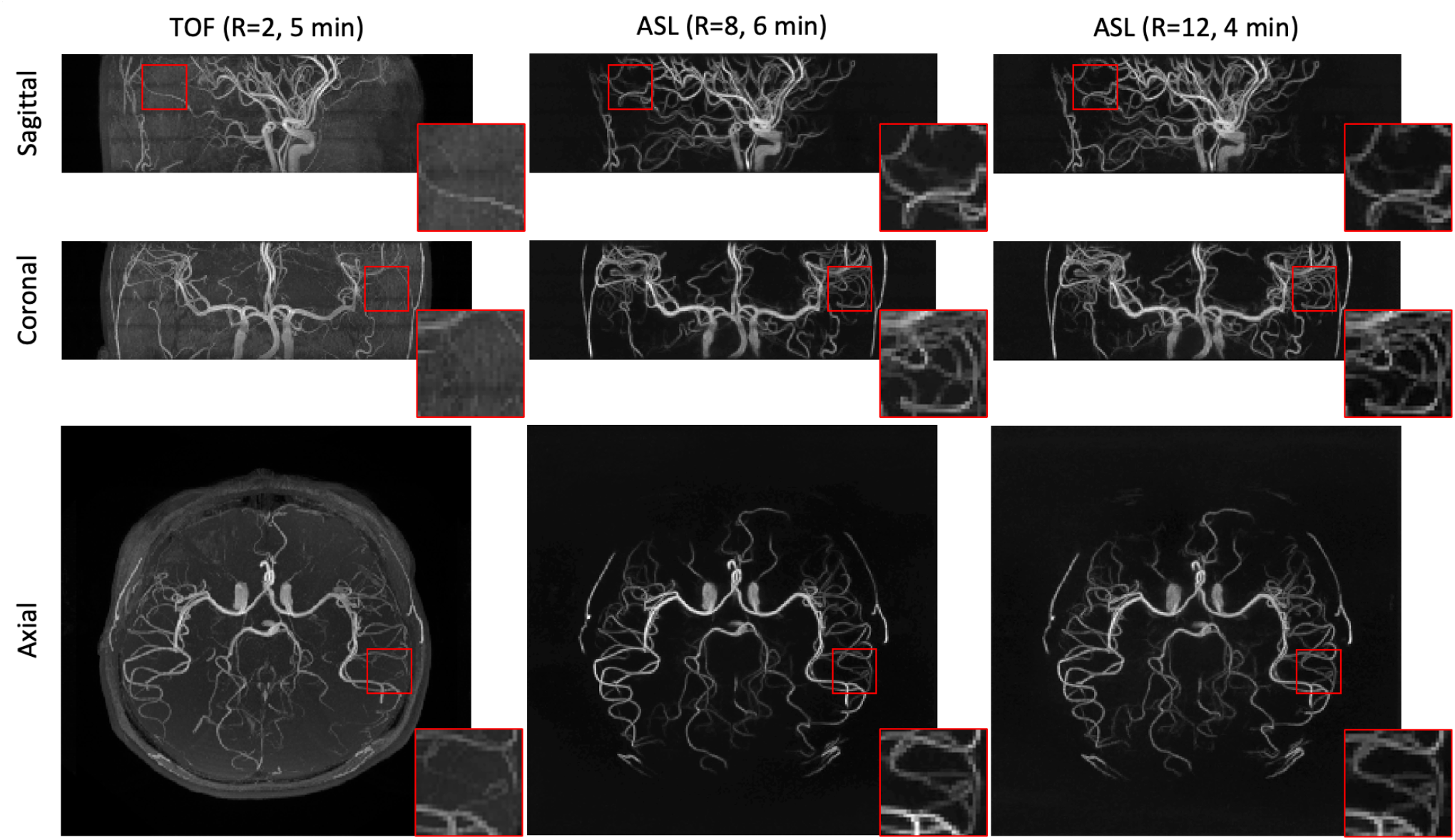
Comparison of multi-slab 2× accelerated 3D TOF-MRA (5 min 4 s) and 4D ASL-MRA acquired with 8× acceleration (6 min) and 12× acceleration (4 min) for large-coverage angiography under prospective undersampling. Zoomed sections show improved distal vessel visibility for ASL compared with TOF.

## 4 Discussion

This study demonstrates that a simulation-driven, few-shot transfer learning framework enables accurate and rapid reconstruction of highly accelerated 4D ASL-MRA, despite the limited availability of fully sampled training data. The proposed method consistently outperformed conventional CS and LLR reconstructions across both retrospective and prospective experiments, with substantial improvements in vessel depiction, temporal fidelity, and computational efficiency. Collectively, these findings support the feasibility of large–coverage, non-contrast dynamic angiography within clinically practical scan times (4–6 min) and reconstruction times (∼15 sec).

Compared with CS and LLR, the proposed method provided superior preservation of vascular structures, particularly in the fine distal vessels. Conventional methods rely on sparsity or low-rank priors that may be less effective for preserving fine vascular detail and variations in signal evolution across the image under high acceleration factors. This limitation is reflected in the blurring of major vessels (e.g., MCA) and loss of small branches in CS and LLR reconstructions, as well as in the structured residuals observed in error maps. In contrast, the proposed method reduces these structured errors and produces more noise-like residuals, as indicated by the higher error entropy, suggesting improved reconstruction of localized vessel signals.

These advantages are maintained under realistic acquisition conditions. In the prospective experiments, the proposed method preserves vessel continuity and achieves superior noise suppression compared with CS and LLR reconstructions. The improved agreement with the reference ASL signal dynamics further indicates that the proposed approach more accurately captures the underlying temporal signal evolution than conventional methods. Results from non-overlapping multi-slab acquisitions demonstrate generalization to extended spatial coverage. Although mild discontinuities were observed at slab boundaries, these artifacts were mitigated in the fully multi-slab acquisition with slab overlap and oversampling.

The underlying reason for these improvements is the combination of simulation-driven pre-training and efficient temporal modeling. Pre-training on large-scale 3D TOF-MRA data provides a strong spatial prior for ASL angiography. The pre-trained model without 4D fine-tuning already exceeded the performance of CS and LLR, indicating that vascular morphology learned from TOF data transfers effectively to ASL-MRA. However, the consistent improvements observed after 4D fine-tuning, particularly in preserving small vessels, suppressing noise, and maintaining temporal signal fidelity, demonstrate that spatial priors alone are insufficient to fully capture the dynamic characteristics of ASL signals. The closer agreement with reference temporal signal curves, further confirms that incorporating temporal information is essential for accurately reconstructing dynamic ASL angiograms.

The proposed lightweight 4D extension enables this temporal modelling efficiently. By retaining the 3D convolutional kernels and introducing temporal fusion modules, the model captures spatiotemporal correlations without substantially increasing the model size and data requirements associated with fully 4D architectures. While the pre-trained model already provided high-quality reconstructions, fine-tuning with 4D extensions using only two experimentally acquired 4D datasets gave substantial further improvements. This highlights the potential practical adaptability of the proposed framework, where the pre-trained model can be efficiently adapted to new scanners, acquisition protocols, or hardware configurations through time- and data-efficient fine-tuning, avoiding the need to collect a large site-specific training dataset.

The ablation studies further clarify the contribution of individual components. Removal of either the histogram-weighted L1+SSIM loss or the simulation-driven pre-training led to consistent degradation in performance, confirming the importance of both elements. Evaluation on time-reversed datasets showed only a modest reduction in performance, suggesting that the model retains reasonable generalizability to substantial change in temporal signal evolution, despite being trained on a fixed temporal ordering. In addition, increasing the number of fine-tuning datasets yields substantial improvements from zero to two datasets, with only marginal gains thereafter, supporting the data-efficient design of the proposed framework.

The comparison with 3D TOF-MRA provides additional context relative to a commonly used angiographic approach. While 3D multi-slab TOF remains a standard technique for structural angiography, it lacks temporal information and may have reduced sensitivity to distal or slow-flow vessels.^55^ The accelerated multi-slab 4D ASL-MRA demonstrated here provides time-resolved angiographic information with improved background suppression, enabling better visualization of distal vessels. Notably, this was achieved with comparable or shorter scan times. Increasing acceleration factor from 8× to 12× introduced only minor image degradation, suggesting that further reductions in scan time may be feasible without compromising the depiction of major vascular structures.

Despite these strengths, several limitations should be considered. First, although the proposed method significantly outperformed conventional approaches, some very small vessels remain challenging to recover. This may reflect residual domain differences between simulated and experimentally acquired data, which could be mitigated through further refinement of the simulation pipeline or incorporation of more diverse training data. Second, validation was performed on a limited cohort of healthy subjects in this work. Further evaluation, including radiological assessment in larger and more diverse patient populations, is required to further establish clinical utility. Third, although the method demonstrated reasonable generalizability, additional and more diverse 4D training data may be needed to better capture atypical or pathological flow patterns. Fourth, this study focused on Cartesian acquisitions with a limited number of temporal frames to enable quantitative evaluation using fully sampled references. While this allowed high quality imaging with large coverage in reasonable scan times, future work will extend the proposed framework to non-Cartesian trajectories to explore the potential for improvements in spatial coverage, temporal resolution, and scan efficiency. Finally, integration into an inline reconstruction pipeline will be necessary to facilitate clinical deployment of the proposed method.

In conclusion, this work demonstrates that combining simulation-driven pre-training with few-shot fine-tuning enables high-quality, rapid reconstruction of highly accelerated 4D ASL-MRA in data-limited scenarios. By leveraging publicly available 3D TOF data and introducing an efficient 4D temporal fusion extension, the proposed framework provides a practical and efficient pathway toward clinically feasible accelerated dynamic ASL angiography.

## Acknowledgments

The Oxford University Centre for Integrative Neuroimaging is supported by core funding from the Wellcome Trust (203139/Z/16/Z and 203139/A/16/Z) with additional support from the NIHR Oxford Biomedical Research Centre (NIHR203311) and the Oxford Health Biomedical Research Centre (NIHR203316). H.L. is supported by the Medical Research Council, the Nuffield Department of Clinical Neurosciences, and Siemens Healthineers through an Oxford-MRC iCASE studentship. M.C. is supported by the Canada Research Chairs Program. P.J. is supported by the Vivensa Foundation and the NIHR Oxford Biomedical Research Centre. T.O. was supported by a Sir Henry Dale Fellowship jointly funded by the Wellcome Trust and the Royal Society (220204/Z/20/Z) and the Podium Institute for Sports Medicine and Technology, University of Oxford.

## Data Availability Statement

The original IXI dataset used for training can be found here: https://brain-development.org/ixi-dataset/. The code underlying the data simulations and model training can be found here: <link to be added upon publication>. Pre-trained models and data underlying the tables and plots in this study can be found here: <link to be added upon publication>. We are currently unable to share the full *in vivo* data due to data protection issues, although the Oxford University Centre for Integrative Neuroimaging is actively working on a solution to this.

