## Supporting Information for "From 3D Time-of-Flight Angiography to Accelerated 4D Arterial Spin Labeling Angiography: A Fast Few-Shot Transfer Learning Approach"

### Supporting Information for “From 3D TOF to 4D ASL: A Fast Few-Shot Transfer Learning Approach for Accelerated ASL Angiography”

Table S1. Parameters for phase simulation. Values for $\sigma_{n_j}$ represent ratios relative to the maximum intensity, while values for $\sigma_{G_j}$ represent ratios relative to the size of the dimensions. $\mathcal{U}(a, b)$ denotes a uniform distribution within a range $[a, b]$ from which the parameters are randomly sampled.

| $j$ | $\sigma_{n_j,vessel}$ | $\sigma_{G_j,vessel}$ | $\sigma_{n_j,background}$ | $\sigma_{G_j,background}$ |
|---|---|---|---|---|
| 1 | $\mathcal{U}$(1.0, 3.0) | $\mathcal{U}$(0.01, 0.05) | $\mathcal{U}$(0.1, 0.25) | 1.0 |
| 2 | $\mathcal{U}$(0.0001, 0.001) | 1.0 | | |

Table S2. Quantitative comparison of reconstruction performance on 8× prospectively accelerated 4D ASL-MRA data. Results presented as mean ± standard deviation across testing datasets. The top-performing method for each metric is highlighted in bold font. The proposed method showed significantly higher performance than all the comparison methods ($p < 0.01$).

| Metric | CS | LLR | Proposed w/o 4D F-T | Proposed |
|---|---|---|---|---|
| PSNR ↑ | 19.30 ± 3.62 | 20.77 ± 0.68 | 22.54 ± 0.98 | **23.57 ± 0.81** |
| SSIM ↑ | 0.2597 ± 0.0721 | 0.2753 ± 0.0283 | 0.3815 ± 0.0447 | **0.4047 ± 0.0459** |
| VM-PSNR ↑ | 25.53 ± 2.20 | 26.37 ± 2.88 | 28.84 ± 1.23 | **30.47 ± 1.21** |
| VM-SSIM ↑ | 0.8972 ± 0.0883 | 0.8968 ± 0.0897 | 0.9270 ± 0.0576 | **0.9395 ± 0.0631** |
| EE ↑ | 2.871 ± 0.153 | 2.914 ± 0.116 | 3.096 ± 0.222 | **3.195 ± 0.195** |

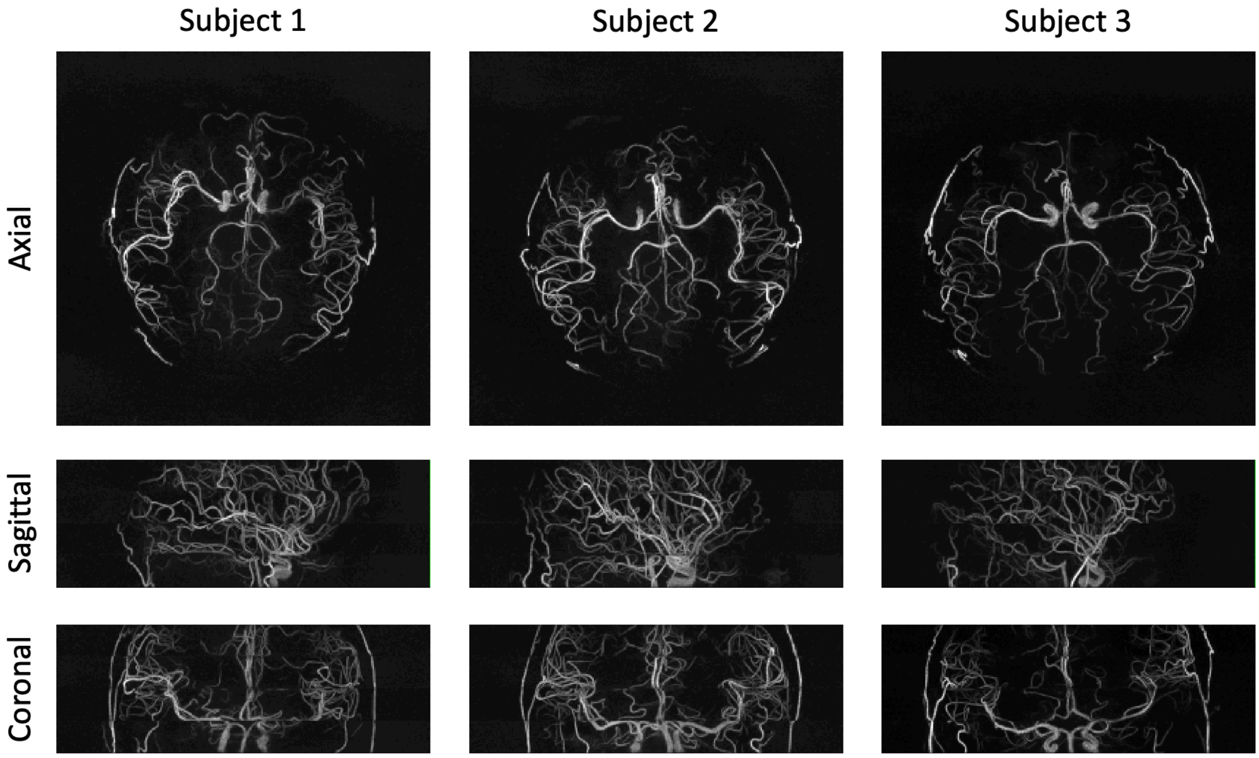


Figure S1. Non-overlapping multi-slab reconstruction for prospectively 8× accelerated 4D ASL-MRA. Due to the non-overlapping acquisition without oversampling, mild discontinuities and boundary artifacts are observed at slab interfaces.

| Metric | Proposed w/o HistLoss | Proposed w/o Pre-Training | Proposed w/o 4D F-T | Proposed | Proposed (Time-Reversed) |
|---|---|---|---|---|---|
| PSNR ↑ | 21.65 ± 0.67 | 21.29 ± 0.84 | 22.54 ± 0.98 | **23.57 ± 0.81** | 22.70 ± 0.66 |
| SSIM ↑ | 0.3355 ± 0.0347 | 0.3457 ± 0.0430 | 0.3815 ± 0.0447 | **0.4047 ± 0.0459** | 0.3909 ± 0.0440 |
| VM-PSNR ↑ | 25.97 ± 0.53 | 26.66 ± 2.76 | 28.84 ± 1.23 | **30.47 ± 1.21** | 29.51 ± 0.80 |
| VM-SSIM ↑ | 0.8856 ± 0.0622 | 0.8976 ± 0.0949 | 0.9270 ± 0.0576 | **0.9395 ± 0.0631** | 0.9328 ± 0.0623 |
| EE ↑ | 2.932 ± 0.074 | 2.995 ± 0.102 | 3.096 ± 0.222 | **3.195 ± 0.195** | 3.139 ± 0.200 |

Table S3. Quantitative results of ablation studies on 8× prospectively accelerated 4D ASL-MRA data. The proposed method showed significantly higher performance than all the comparison methods ($p < 0.05$). The table format follows Table S2.

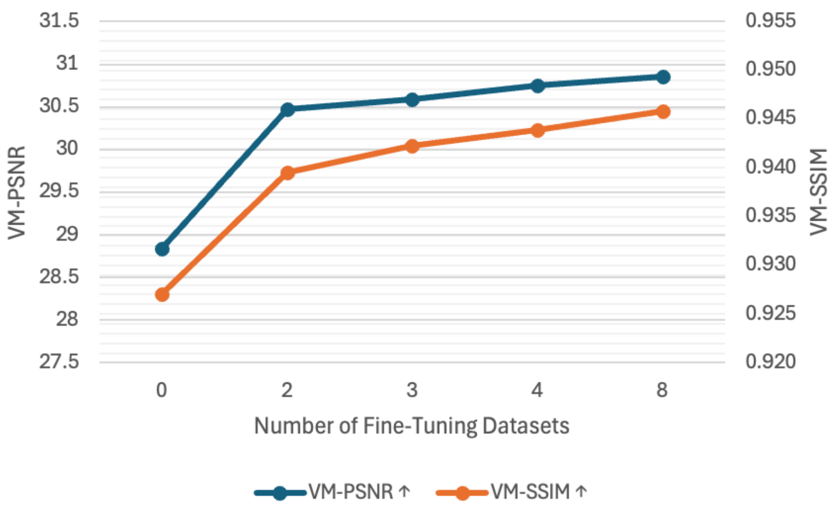


Figure S2. Quantitative evaluation of reconstruction performance as a function of the number of fine-tuning datasets.
